# Customized Protein Nanoreactors for Affibody-Directed Activation of 5-Fluorcytosin in HER2-Positive Cells

**DOI:** 10.1101/2025.09.11.675605

**Authors:** Mariia Zmyslia, Martin Holzer, Claudia Jessen-Trefzer

## Abstract

The development of targeted therapies for HER2-positive cancers remains critical due to resistance and toxicity challenges in current treatments. Here, we present the rational engineering of protein-based encapsulin nanocompartments for selective catalytic prodrug activation. Encapsulins provide precise cargo loading, exceptional stability, and versatile engineerability, making them ideal nanoreactors for therapeutic applications. Our encapsulin constructs encapsulate tandem cytosine deaminase enzymes and display HER2-specific affibodies on their exterior, enabling precise cellular targeting. These engineered nanoreactors catalyze the efficient conversion of the prodrug 5-fluorocytosine (5-FC) into the cytotoxic agent 5-fluorouracil (5-FU), yielding an 83% reduction in viability of HER2-overexpressing SKOV3 cells. Structural characterization using native gel electrophoresis confirms stable assembly with functional affibody presentation. This enzyme-prodrug approach showcases how supramolecular protein architectures can serve as customizable platforms for affibody-directed, enzyme-mediated therapy, offering a promising strategy to enhance therapeutic specificity and minimize systemic side effects in HER2-positive cancer treatment.

**Significance Statement:** This work establishes encapsulins as a new class of programmable therapeutic nanoreactors by integrating selective affibody-mediated targeting with enzyme-prodrug catalysis in HER2-positive cancer cells. Unlike virus-like particles, which often lack precise cargo loading and structural robustness, encapsulins enable dual engineering of both interior and exterior domains for stable, multifunctional assemblies. Compared to antibody–drug conjugates, these nanoreactors achieve amplified drug generation through localized prodrug activation, overcoming payload limitations and reducing systemic toxicity. This platform introduces a versatile supramolecular strategy for targeted cancer therapy that exceeds current delivery technologies.

## INTRODUCTION

Human Epidermal Growth Factor Receptor 2 (HER2) is a critical member of the epidermal growth factor receptor (EGFR) family involved in the regulation of cell growth, proliferation and differentiation.^1^ HER2 is overexpressed in a subset of cancers,^2–5^ particularly breast and ovarian cancer, which significantly impacts prognosis and treatment strategies.^6,7^ Studies indicate that patients with HER2-positive tumors experience poorer outcomes due to increased invasiveness, higher recurrence rates, and a greater propensity for metastasis compared to HER2-negative cases.^6,8^ In clinical practice, the identification of HER2 overexpression plays a crucial role in guiding therapeutic decisions. Trastuzumab (Herceptin), a monoclonal antibody that specifically targets the HER2 receptor, has been pivotal in the management of HER2-positive breast cancer. Its use has significantly improved patient outcomes by inhibiting HER2-mediated signaling pathways and promoting antibody-dependent cellular cytotoxicity.^9^ However, resistance to trastuzumab is a significant clinical challenge, as some patients experience primary resistance while others develop secondary resistance during treatment. This necessitates the exploration of alternative HER2-targeted strategies that may complement or expand the existing therapeutic landscape.

One innovative strategy to enhance targeted cancer therapy involves the covalent attachment of cytotoxic drug molecules to monoclonal antibodies or alternative recognition elements, creating antibody-drug conjugates (ADCs).^10,11^ While ADCs have demonstrated clinical promise, their drug loading capacity is inherently limited by the relatively low number of conjugation sites per antibody molecule.^12,13^ To circumvent this constraint, enzyme-prodrug approaches, where catalytic enzymes are selectively delivered to tumor tissue, allow for local conversion of benign prodrugs into highly potent cytotoxic agents, thereby amplifying the therapeutic effect at the target site and reducing systemic toxicity.^14^ Cancer-targeted enzymes can convert a prodrug into a greater quantity of active drug molecules at the tumor site, thereby maximizing therapeutic efficacy and potentially minimizing the systemic toxicity associated with traditional chemotherapy. However, a systemic application of enzymes presents challenges, particularly regarding enzyme stability and the immunogenicity of non-human enzymes used in Antibody-Directed Enzyme Prodrug Therapy (ADEPT). Encapsulation of enzymes within robust nanoscale carriers, such as protein-based encapsulins, offers a promising solution by providing a protective microenvironment for the enzyme catalyst, while enabling further functionalization for targeted delivery.^15,16^

Encapsulins are prokaryotic protein-based nanocompartments widely distributed among bacterial and archaeal phyla, typically measuring 24–42 nm in diameter and assembling from 60–240 subunits in an icosahedral arrangement.^17,18^ They self-assemble from shell proteins that adopt the viral HK97 fold, forming a robust and stable scaffold that encases specialized cargo proteins.^19^ In native systems, these cargo molecules – commonly ferritin-like proteins, peroxidases, or desulfurases – are recruited through a short C-terminal encapsulin localization sequence (ELS) or an alternative N-terminal targeting motif.^20–22^ This encapsulation shields cargo proteins from proteolysis, thermal degradation, or pH fluctuations.^18,21,23,24^ Phylogenetic analyses have revealed four distinct encapsulin families (type I–IV), each characterized by unique sequence features, structural variations, and operon organizations.^25^

Studies show that purified encapsulin derived from *Thermotoga maritima* offers several key advantages, including colloidal stability, storability, and compatibility with blood when administered intravenously in animal models. Furthermore, it demonstrates an excellent nanosafety profile in mice, with no significant weight loss or adverse pathology observed.^26^ Like other intravenously applied protein nanocages (PNCs), encapsulins can induce the production of PNC-specific antibodies, which play a key role in their clearance from the body. Enhancing the therapeutic viability of these nanocages can be achieved by using site-directed mutagenesis to modify their immunogenicity, specifically by removing certain B- and T-cell epitopes.^27^ Encapsulin, in particular, shows a remarkable ability to tolerate various amino acid modifications, offering a flexible platform for engineering nanocarriers with improved biochemical properties and reduced immunogenicity.^20,28–33^

Beyond this, the exceptional amenability of encapsulins to genetic engineering allows researchers to modify both the interior and exterior of the protein shell for diverse biotechnological and therapeutic applications.^18,19^ By incorporating alternative cargo proteins through an ELS, encapsulins can be repurposed to compartmentalize enzymes employed in ADEPT, providing a protective environment that enhances enzyme stability and function. In parallel, their exterior surface can be functionalized with targeting moieties to enable selective interaction with defined cell types, offering a route to localized prodrug activation. This dual-modification capability positions encapsulins as versatile scaffolds for developing targeted enzyme delivery systems.

Among the various targeting moieties, affibodies stand out for their compact size, high target affinity, and efficient bacterial expression, making them ideal for functionalizing protein nanocages to achieve tumor specificity. Derived from the IgG-binding domains of Staphylococcal protein A, affibodies are small (∼58 amino acids) proteins arranged in a stable three-helix bundle.^34^ HER2-specific affibodies have been explored for targeted delivery of small drug molecules,^35,36^ photosensitizers,^37^ imaging reagents,^38–40^ enzymes,^41,42^ and organometallic catalysts,^43^ as well as for the delivery of nanoparticles and other synthetic carriers.^30,44–46^

In light of these developments, we have chosen to engineer an encapsulin platform for affibody-directed enzyme-prodrug activation in HER2-positive cell models. This strategy centers on the encapsulation of yeast cytosine deaminase for localized conversion of the prodrug 5-fluorocytosine into 5-fluorouracil, a cytotoxic agent widely used in chemotherapy. This work pioneers a new therapeutic class that merges the programmability of protein nanocages with the selectivity of affibody targeting and the amplification power of enzyme-prodrug therapy.

## MATERIALS AND METHODS

Please refer to the supporting information for detailed descriptions of materials, procedures, complete protein sequences, and additional supporting figures.

## RESULTS AND DISCUSSION

### Construction of the encapsulated cytosine deaminase Enc{tdCD}

We introduced the encapsulin gene MSMEI_5672 from *Mycobacterium smegmatis* mc^2^155, tagged with a C-terminal Strep-tag, under the first T7 promoter of the pETDuet-1 vector. Given that yeast cytosine deaminase (CD) functions as an obligate homodimer, and our previous findings have shown that successful dimer formation and retention of enzymatic activity upon encapsulation require the enzyme monomers to be connected by a flexible linker^47^, we cloned CD as a tandem enzyme (tdCD) under the second T7 promoter of pETDuet-1 vector. This was achieved by linking two copies of the CD gene FCY1 with a flexible linker sequence and appending a C-terminal ELS sequence for directing tdCD to the interior of encapsulin (as illustrated in Figure 1A; for PCR primers see Table S1). Following transformation into *E. coli*, we purified the encapsulin particles using affinity chromatography, targeting the surface-exposed Strep-tag at the C-terminus of the encapsulin shell. Subsequently, encapsulin was further polished through size-exclusion chromatography (Figure 1D). As a control, we cloned and produced encapsulin harboring eGFP, following the same purification procedure described above.

**Figure 1.**
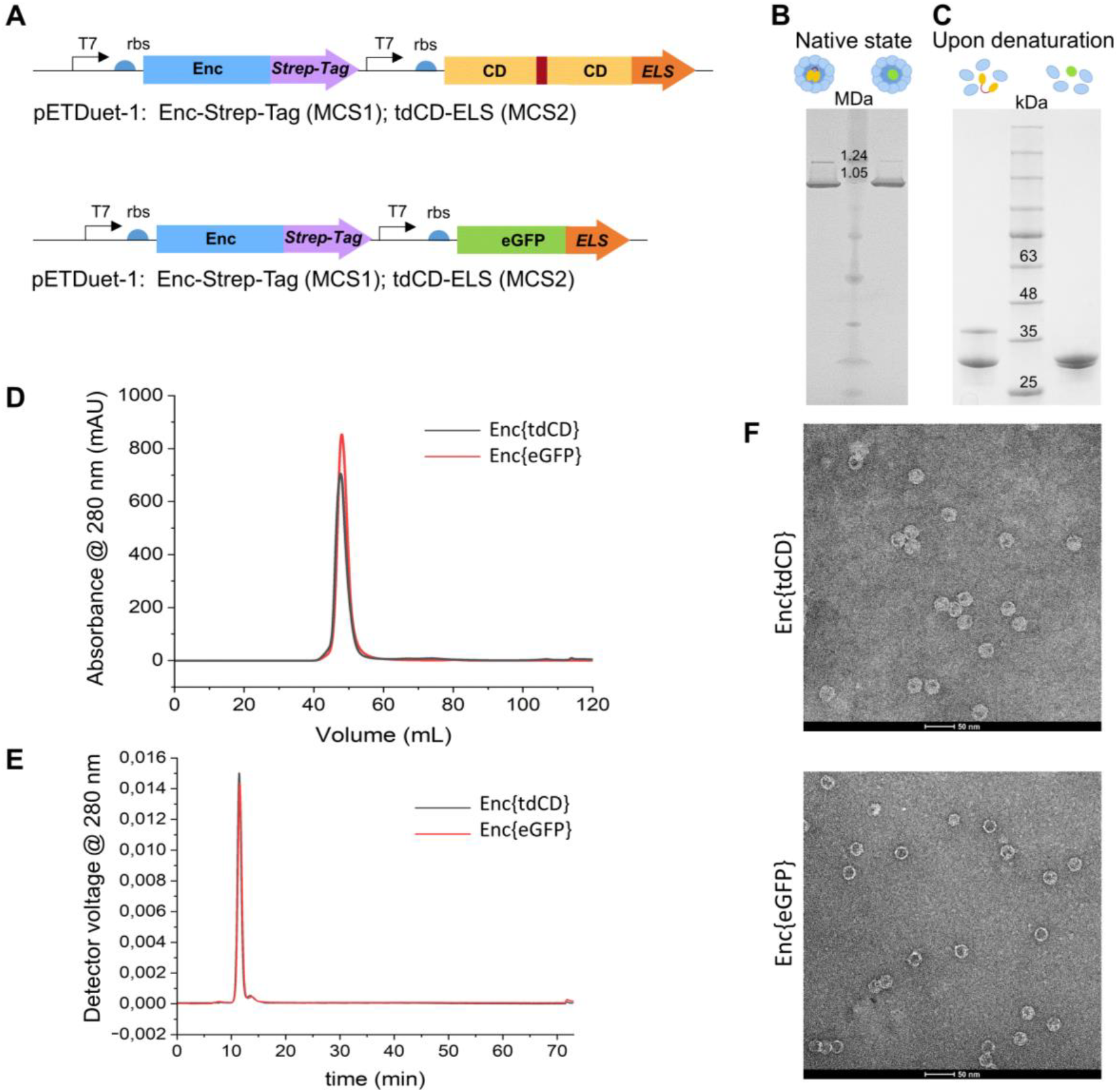
Design and characterization of Enc{tdCD} and Enc{eGFP}. A) Schematic outline of the cloned constructs. B) BN-PAGE and C) SDS-PAGE analysis after purification via affinity chromatography on a StrepTrap HP column, followed by size exclusion chromatography on a HiLoad 16/600 Superdex 200 PG column. D) Representative chromatogram of a size-exclusion column chromatography run (HiLoad 16/600 Superdex 200 PG column). E) AF4 fractionation profile using UV (280 nm). F) TEM analysis of Enc{tdCD} and Enc{eGFP}. Bar: 50 nm.

### Characterization of encapsulin constructs Enc{tdCD} and Enc{eGFP}

The structural integrity and oligomeric assembly of the purified encapsulin nanocompartments were analyzed using Blue Native Polyacrylamide Gel Electrophoresis (BN-PAGE), revealing large supramolecular protein complexes (Figure 1B). The dominant lower band corresponds to the canonical T1 encapsulin capsids, while a fainter upper band likely represents higher-order assemblies or aggregates. Migration patterns relative to protein standards suggested observed sizes of approximately 1.2 and 0.9 MDa, respectively, both lower than the theoretical molecular mass of the full 60-mer capsid (1.8 MDa). This anomalous migration mirrors previous reports for protein nanocages, where potentially molecular shape and intrinsic charge influence gel mobility under native conditions.^24,47,48^

Under denaturing conditions, SDS-PAGE revealed two distinct protein bands corresponding to encapsulin monomers (30 kDa) and either tandem cytosine deaminase (38.4 kDa) or eGFP (28.3 kDa) in the Enc{tdCD} and Enc{eGFP} constructs, respectively (Figure 1C).

To achieve high-resolution characterization and distinguish properly assembled particles from potential aggregates or defective structures, we employed AF4-MALS, a state-of-the-art method for virus-like particle (VLP) characterization. AF4-MALS analysis identified a primary, well-resolved peak at 12 mL, corresponding to correctly assembled T1 capsids, and a minor aggregation peak at 15 mL (Figure 1E), contributing to less than 5% of the total area under the curve (AUC). The observation of two distinct peaks in AF4-MALS corresponds well with the two bands observed in BN-PAGE. The mass-weighted mean of the assigned molecular masses for the primary peaks were 2513 ± 14 kDa for Enc{tdCD}, 2284 ± 28 kDa for Enc{eGFP}, and 1852 ± 24 kDa for empty encapsulin shell. From these data, we estimated the cargo load per nanocompartment to comprise 18.7 ± 0.2 tdCD molecules for Enc{tdCD} and 17.0 ± 1.0 eGFP molecules for Enc{eGFP}.

To further confirm the formation of encapsulin nanocompartments, Transmission Electron Microscopy (TEM) and Dynamic Light Scattering (DLS) analyses were performed (Table 1). TEM micrographs demonstrated well-defined, spherical nanostructures with an average outer diameter of 25.0 ± 1.3 nm (n = 100) for Enc{tdCD} and 22.4 ± 1.6 nm (n = 100) for Enc{eGFP} (Figure 1F). DLS measurements revealed hydrodynamic diameters of 28.8 ± 0.1 nm (PDI = 0.11 ± 0.01) for Enc{tdCD} and 29.5 ± 0.3 nm (PDI = 0.14 ± 0.01) for Enc{eGFP} (Table 1, Figure S1), further confirming the formation of stable nanocompartments.

**Table 1:** Size characterization of encapsulin constructs via TEM, AF4-MALS, and DLS.

| Sample | TEM external diameter (nm) | AF4-MALS diameter of gyration (nm) | DLS hydrodynamic diameter (nm) |
| --- | --- | --- | --- |
| Enc | $24.0 \pm 2.3$ | $20.2 \pm 0.3$ | $29.3 \pm 0.1$ |
| Enc{tdCD} | $25.0 \pm 1.3$ | $19.2 \pm 0.0$ | $28.8 \pm 0.1$ |
| Enc{eGFP} | $22.4 \pm 1.6$ | $18.4 \pm 1.2$ | $29.5 \pm 0.3$ |
| <sup>ZHER</sup>Enc{tdCD} | $27.6 \pm 2.0$ | $21.2 \pm 0.4$ | $29.6 \pm 0.0$ |
| <sup>ZHER</sup>Enc{eGFP} | $26.2 \pm 1.3$ | $21.0 \pm 0.6$ | $28.7 \pm 0.1$ |

These results collectively validate the successful encapsulation and structural integrity of the engineered nanocompartments, paving the way for their downstream applications.

### In vitro and in cellulo prodrug activation by Enc{tdCD}

To verify that the encapsulated cytosine deaminase (CD) retains its enzymatic activity, an in vitro assay was performed by incubating Enc{tdCD} with the substrate 5-FC. HPLC analysis confirmed the conversion of 5-FC to the product 5-FU within one hour of incubation (Figure 2A, Figure S2). These results indicate that the encapsulation process preserves the enzymatic activity of cytosine deaminase. Building on the in vitro results, we next evaluated whether Enc{tdCD} retains its enzymatic activity in living cells, aiming to explore its potential in cell-based prodrug activation models. Specifically, we examined its capacity to convert the prodrug 5-FC into 5-FU, a cytotoxic compound widely used in chemotherapy, by assessing cell viability. To investigate this, Enc{tdCD} was incubated with the mouse monocyte cell line J774A.1, known to internalize native encapsulin nanocompartments.^49,50^ Following incubation, unbound nanoparticles were removed by washing, and the 5-FC prodrug was subsequently added. This treatment resulted in a significant reduction in cell viability to 18.1% (Figure 2B) compared to controls treated with Enc{eGFP}, indicating effective intracellular prodrug conversion mediated by Enc{tdCD}. Importantly, neither Enc{tdCD} nor Enc{eGFP} exhibited inherent toxicity to the cells when applied alone; in fact, they resulted in small increases in cell viability (12.5% and 7.9%, respectively).

**Figure 2.**
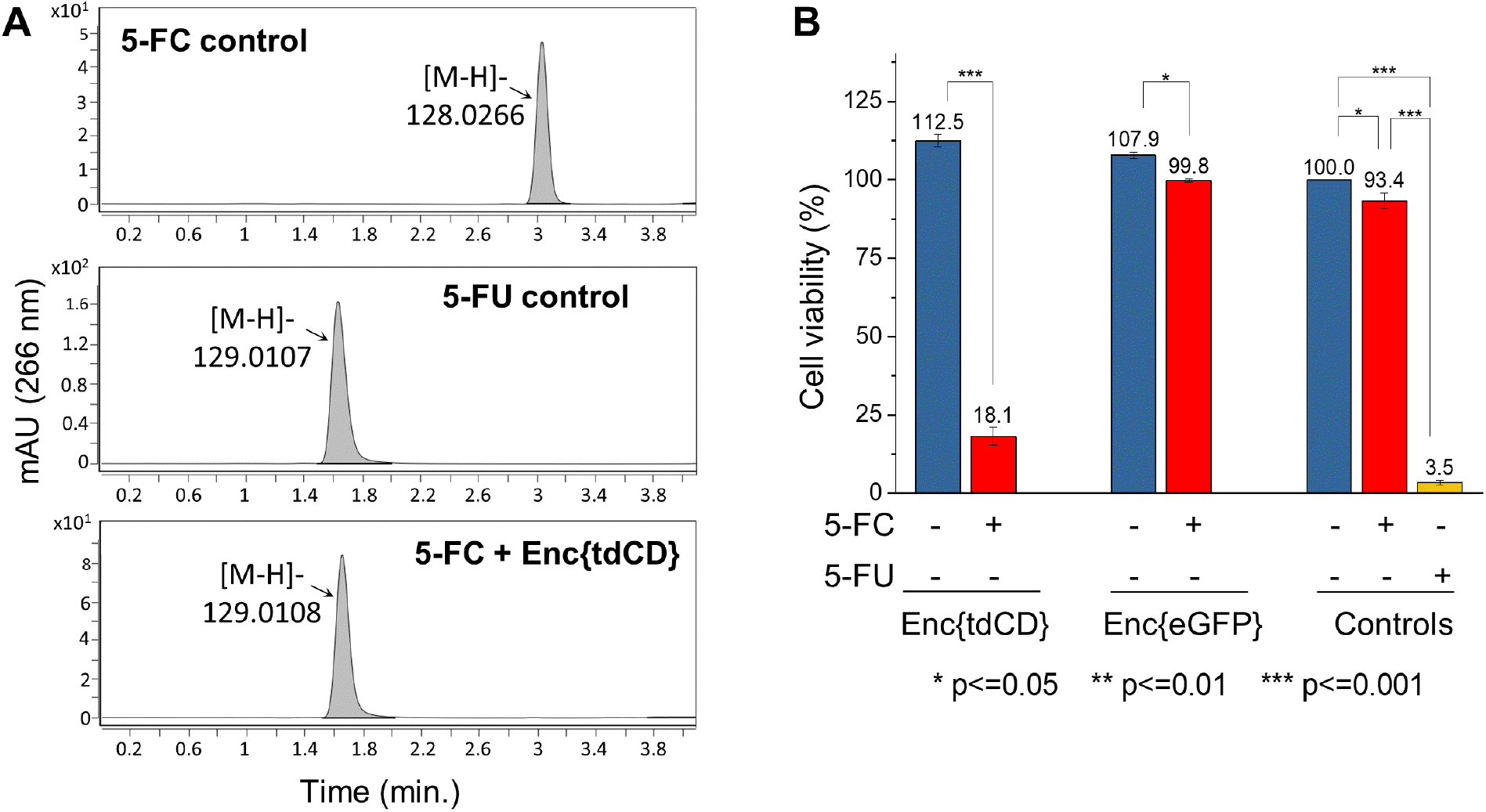
Prodrug activation by Enc{tdCD}. A) HPLC analysis of in vitro Enc{tdCD}-mediated conversion of 5-fluorocytosine (5-FC) to 5-fluorouracil (5-FU). B) Impact of in cellulo prodrug activation by Enc{tdCD} on J774A.1 cell viability assessed by MTS assay.

### Evolution of encapsulin shell for specific cell targeting

These promising results motivated us to design surface modification of the encapsulin shell for targeted delivery to specific cell types. We selected the HER2 receptor, whose abnormal expression is correlated with the development and progression of various cancers,^2–5^ as well as enhanced invasiveness and resistance to chemotherapy.^6,7^ In recent years, a variety of affibodies targeting the HER2 receptor have been reported.^51,52^ Affibodies are antibody mimetics of small size, high target affinity and ease of bacterial expression. Among these, the ZHER_2:342_ affibody demonstrates strong HER2 binding (22 pM)^52^ and retains its specificity even when fused at its N- or C-terminus, making it an ideal candidate for decorating the surface of the encapsulin shell.^53,54^

Initially, we constructed a dual-expression vector analogous to the one for Enc{tdCD} by placing an encapsulin-ZHER_2:342_ affibody fusion (with a C-terminal His-tag) under the first T7 promoter in the pETDuet-1 vector and the tdCD sequence under the second T7 promoter. Since C-terminal encapsulin fusions, such as purification tags, have been shown to be presented on the outer surface of the encapsulin shell,^24,31,47^ we anticipated that the affibody would likewise be presented on the surface. However, this single-vector design failed to yield fully assembled nanocompartments (^Full-ZHER^Enc{tdCD}), as indicated by the absence of high-molecular-weight species in BN-PAGE (Figure S3), likely due to steric hindrance introduced by the bulky affibody moiety, which disrupted proper capsid assembly.

To overcome this limitation, we adopted a co-expression strategy in which a wild-type encapsulin gene was expressed alongside the affibody-fused variant and tdCD acrgo, ensuring that mixed populations of monomers – both with and without the affibody – are available for assembly. This strategy mitigates steric interference and facilitates the successful formation of stable nanocompartments. Accordingly, a set of plasmids was prepared for the co-expression of three proteins required for the assembly of the nanocompartment (Figure 3*Figure 2A*). The sequence encoding the encapsulin-ZHER_2:342_ affibody fusion, followed by C-terminal His-tag was cloned under the first T7 promoter of the pCDFDuet-1 vector. The unmodified encapsulin gene sequence was inserted under the second T7 promoter of the same vector. The pETDuet-1 vector was designed to accommodate the sequence for the cargo protein, either tdCD or eGFP followed by a C-terminal ELS sequence. Each combination of plasmids was co-transformed into *E. coli* BL21 Star (DE3), and following expression, the fully assembled nanocompartments ^ZHER^Enc{tdCD} and ^ZHER^Enc{eGFP}, were purified via affinity chromatography and further polished through size-exclusion chromatography (Figure 3D).

**Figure 3.**
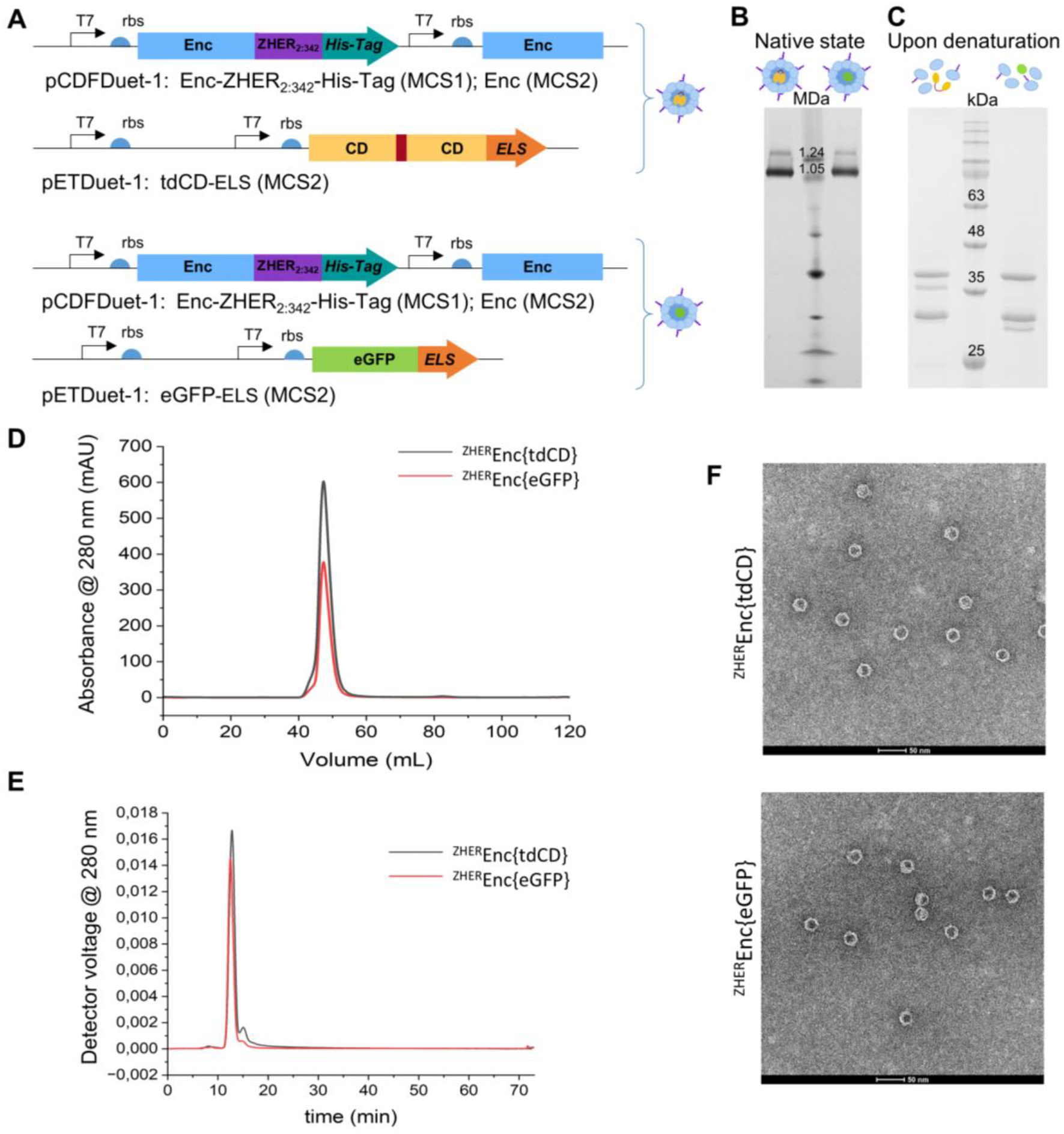
Design and characterization of ^ZHER^Enc{tdCD} and ^ZHER^Enc{eGFP}. A) Schematic outline of the cloned constructs. B) BN-PAGE and C) SDS-PAGE analysis after purification via affinity chromatography on a HisTrap HP column, followed by size exclusion chromatography on a HiLoad 16/600 Superdex 200 PG column. D) Representative chromatogram of a size-exclusion column chromatography run (HiLoad 16/600 Superdex 200 PG column). E) AF4 fractionation profile using UV (280 nm). F) TEM analysis of ^ZHER^Enc{tdCD} and ^ZHER^Enc{eGFP}. Bar: 50 nm.

### Characterization of shell-modified encapsulin constructs ^ZHER^Enc{tdCD} and ^ZHER^Enc{eGFP}

For encapsulin constructs with a ZHER_2:342_ affibody-decorated shell surface, BN-PAGE revealed two bands analogous to previous constructs: a strong lower band and a weaker upper band, corresponding to approximate sizes of 1.3 and 1.1 MDa, respectively (Figure 3*Figure 2B*). Compared to the unmodified constructs, these results indicate an increase in molecular weight, consistent with the successful integration of the affibody into the encapsulin shell. The broader and less distinct appearance of these bands can be potentially attributed to the heterogeneity introduced during assembly. The encapsulin shell comprises both wild-type and affibody-fused encapsulin monomers. Wild-type encapsulin plays a crucial role in preventing steric hindrance between affibody-fused monomers during the assembly process. As the ratio of these two protein types is not fixed and may vary slightly, this variability likely contributes to the observed band broadening.

Under denaturing SDS-PAGE conditions, three distinct protein bands were observed: encapsulin monomers (28.8 kDa), affibody-fused encapsulin monomers (37.8 kDa), and either tandem cytosine deaminase (38.4 kDa) or eGFP (28.3 kDa) in the ^ZHER^Enc{tdCD} and ^ZHER^Enc{eGFP} constructs, respectively (Figure 3C).

To quantify the stoichiometry of affibody display, we performed densitometric analysis on the SDS-PAGE gels (Figure S4). Given that each encapsulin shell comprises 60 self-assembled protomers, this quantitative analysis enabled precise estimation of affibody fusion density. Results indicate that shells of ^ZHER^Enc{tdCD} contain approximately 21.0 ± 0.8 affibody-modified and 39.0 ± 0.8 unmodified subunits, while ^ZHER^Enc{eGFP} constructs incorporate 22.2 ± 0.4 ^ZHER^Enc and 37.8 ± 0.4 Enc protomers. These ratios support efficient site-specific incorporation of affibody motifs onto the shell exterior via protein engineering.

AF4-MALS analysis of the affibody-modified constructs showed a primary, well-resolved peak at 12 mL, with a secondary peak at 15 mL (Figure 3E). While the secondary peak was more pronounced compared to the unmodified constructs, the overall profile suggested a substantial degree of sample homogeneity. The increased mass-weighted mean of molecular masses for the primary peaks, determined to be 2574 ± 22 kDa for ^ZHER^Enc {tdCD} and 2376 ± 36 kDa for ^ZHER^Enc {eGFP}, provides further evidence of successful surface modification. Upon nanocompartment assembly, the affibody protrudes from the external surface of the encapsulin, which is expected to increase both the outer diameter and the diameter of gyration. This was confirmed through AF4-MALS and TEM, both of which detected larger sizes for the surface-modified constructs compared to the unmodified ones. AF4-MALS estimated mass-weighted mean diameter of gyration of 21.2 ± 0.4 nm and 21.0 ± 0.6 nm for the ^ZHER^Enc{tdCD} and ^ZHER^Enc{eGFP} constructs, respectively.

TEM micrographs revealed spherical nanostructures with well-preserved morphology and an increased average outer diameters compared to the unmodified constructs: 27.6 ± 2.0 nm (n = 100) for ^ZHER^Enc{tdCD} and 26.2 ± 1.3 nm (n = 100) for ^ZHER^Enc{eGFP} (Figure 3F). DLS measurements further confirmed these findings, revealing consistent hydrodynamic diameters of 29.6 ± 0.0 nm (PDI = 0.18 ± 0.09) for ^ZHER^Enc{tdCD} and 28.7 ± 0.1 nm (PDI = 0.11 ± 0.01) for ^ZHER^Enc{eGFP} (Figure S1, Table 1).

Collectively, these results confirm the successful incorporation of the ZHER_2:342_ affibody into the encapsulin shell while maintaining its structural integrity.

### Cell-based affinity characterization

The binding specificity of the affibody-functionalized encapsulin construct ^ZHER^Enc{eGFP} was evaluated using the HER2-overexpressing SKOV-3 cell line. As a control, the non-modified Enc{eGFP} construct was tested in parallel. For fluorescent microscopy SKOV-3 cells were treated with either ^ZHER^Enc{eGFP}, Enc{eGFP}, or left untreated. Green fluorescence was observed exclusively in cells incubated with ^ZHER^Enc {eGFP} (Figure 4A**Fehler! Verweisquelle konnte nicht gefunden werden**.). These results indicate that only the affibody-decorated construct binds specifically to HER2-positive SKOV-3 cells.

**Figure 4.**
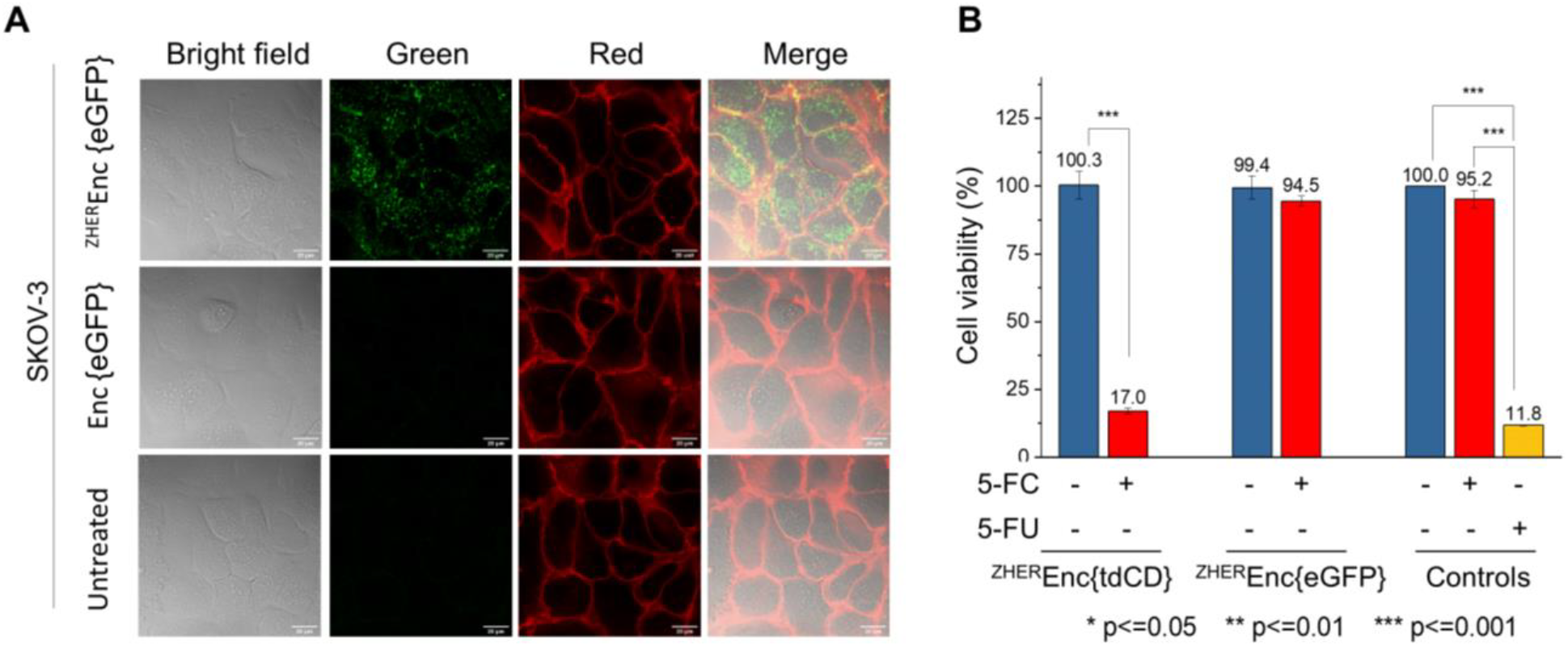
Microscopy analysis and prodrug activation in SKOV-3 cells. A) Fluorescence microscopy and bright-field images of SKOV-3 cells treated for 90 min with either 20 nM ^ZHER^Enc{eGFP}, 20 nM Enc{eGFP}, or left untreated. B) Impact of in cellulo prodrug activation by ^ZHER^Enc{tdCD} on SKOV-3 cell viability assessed by MTS assay.

Flow cytometric analysis was carried out to quantitatively assess binding specificity of eGFP-labeled constructs toward HER2-overexpressing (SKOV-3) and HER2-low (OV7^55,56^) cell lines. Treatment with Enc{eGFP} (lacking affibody) showed negligible cell-associated fluorescence, whereas ^ZHER^Enc{eGFP} produced a pronounced, concentration-dependent increase in mean fluorescence intensity (MFI) and percentage of GFP-positive SKOV-3 cells, reaching up to a 62-fold MFI elevation at 20 pmol per 1 × 106 cells (Figure S5). In OV7 cells, with low endogenous HER2, only minimal increases in MFI and GFP positivity were seen upon ^ZHER^Enc{eGFP} exposure, confirming effective selective targeting via the surface-displayed affibody.

### In cellulo prodrug activation by ^ZHER^Enc{tdCD}

To evaluate the catalytic function of encapsulated cytosine deaminase, we measured the conversion of 5-fluorocytosine (5-FC) to 5-fluorouracil (5-FU) in SKOV-3 cells (Figure 4B). Following cell treatment with ^ZHER^Enc{tdCD}, with or without subsequent addition of 4 mM 5-FC, cell viability assays revealed pronounced cytotoxicity exclusively in wells receiving both encapsulin-based biocatalyst and prodrug, confirming locally triggered prodrug activation propagated by the shell-encased enzyme (Figure 4B). Controls with 5-FU (active agent) or 5-FC (prodrug alone) supported the requirement for enzyme-mediated prodrug conversion. Treatment with the positive control 2 mM 5-FU (higher concentrations could not be tested due to solubility limitations in cell culture medium), led to slightly higher cytotoxicity (11.8%) compared to ^ZHER^Enc{tdCD} with 4 mM 5-FC (17.0%) condition. This difference may be attributed to the encapsulation context, where delayed diffusion of the enzymatically generated 5-FU from the nanocompartment could attenuate its immediate cytotoxic effect.

This outcome demonstrates the prodrug-activating capability of ^ZHER^Enc{tdCD} in SKOV-3 cells. However, several factors may influence the observed efficacy. One key consideration is the efficiency of 5-FC conversion within the encapsulin shell. Previous studies have shown that encapsulin pores function as selective molecular sieves, with transport governed by the size and charge properties of both the pores and the diffusing molecules.^57–59^ Reduced catalytic efficiency in protein-based nanoreactors is often attributed to an incompatibility between the selective permeability of the protein shell and the physicochemical properties of the encapsulated enzyme’s substrate.^59^ In this context, limited diffusion of 5-FC into the nanocompartment may reduce substrate availability for enzymatic conversion, while restricted release of 5-FU from the capsid could delay or diminish its cytotoxic effects.

Future investigations may employ rational design of encapsulin pore size and surface charge through site-directed mutagenesis to modulate molecular flux and enhance substrate/product diffusion, while preserving the structural and catalytic integrity of the protein nanocompartments.^59^ Notably, encapsulins have demonstrated remarkable robustness toward pore-region mutations, providing a versatile supramolecular platform for engineering of chemical selectivity, permeability, and throughput − critical parameters for optimizing catalytic nanoreactor efficiency in therapeutic and synthetic chemistry contexts.^28,29^

Together, these findings demonstrate ^ZHER^Enc{tdCD}-mediated prodrug activation in SKOV-3 cells and open avenues for enhancing efficacy through structural tuning of the nanoreactor.

## CONCLUSION

This study represents the successful application of a designer encapsulin in targeted (affibody-directed) enzyme prodrug therapy. We have demonstrated the encapsulation of tandem cytosine deaminase within an engineered encapsulin nanocompartment, highlighting its potential for targeted prodrug activation in a cellular context, particularly relevant for cancer therapy. Our approach hold the potential to significantly enhance the therapeutic efficacy of 5-FC by facilitating its localized conversion to the active drug 5-FU directly within the microenvironment of HER2-positive cancer cells. By ensuring that enzymes remain concentrated at the site of action this strategy has the potential to drastically reduce systemic toxicity associated with traditional chemotherapy.

## AUTHOR CONTRIBUTIONS

Conceptualization: M.Z. and C.JT.; methodology: M.Z., M.H..; validation: M.Z., M.H., C.JT.; resources: C.JT., M.H.; data curation: M.Z.; writing - original draft preparation: M.Z.; writing - review and editing: M.Z., M.H., C.JT. All authors have read and agreed to the published version of the manuscript.

## ACKNOWLEGEMENTS

We acknowledge support by Ralf Thomann during TEM measurements at the Freiburg Center for Interactive Materials and Bioinspired Technologies (FIT), as well as by Jürgen Beck for technical support. C.JT. acknowledges funding by the Deutsche Forschungsgemeinschaft (DFG, German Research Foundation) under Germany’s Excellence Strategy–EXC-2189. C.JT. and M.Z., acknowledge funding by the Deutsche Forschungsgemeinschaft grant RTG 2202 (Transport across and into membranes). C.JT. acknowledges funding by the Deutsche Forschungsgemeinschaft grant JE 782/4-1 and JE 782/6-1. We furthermore thank all members of LIC for supporting fluorescence imaging efforts and image analysis, funded by DFG-Funding INST 380/109-1 FUGG (Zeiss LSM 880). Open Access funding enabled and organized by Projekt DEAL. We thank the Lighthouse Core Facility, Medical Faculty, University of Freiburg (Project Number 2023/A2-Fol) for assistance with cell sorting. We also thank Dr. Elisabetta Grillo (University of Brescia) for generously providing OV-7 cells and Prof. Tilman Brummer (University of Freiburg) for generously providing the SKOV-3 cell line.

## DATA AVAILABILITY STATEMENT

Data is available within the article or its supplementary materials.

## TOC Text

Engineered encapsulin nanoreactors combine HER2-specific affibody targeting with cytosine deaminase-mediated prodrug activation for selective cancer therapy. These robust protein cages achieve precise cargo loading, amplified local drug generation to reduce systemic toxicity, offering a programmable alternative to virus-like particles and antibody–drug conjugates.

## REFERENCES

1 Yarden Y, Sliwkowski MX. Untangling the ErbB signalling network. Nat Rev Mol Cell Biol. 2001;2(2):127–137. doi:10.1038/35052073.

2 Stigbrand T, Carlsson J, Adams GP, eds. Targeted Radionuclide Tumor Therapy: Biological aspects. Dordrecht: Springer Netherlands; 2008.

3 Morote J, Torres I de, Caceres C, Vallejo C, Schwartz S, Reventos J. Prognostic value of immunohistochemical expression of the c-erbB-2 oncoprotein in metastasic prostate cancer. Int. J. Cancer. 1999;84(4):421–425. doi:10.1002/(SICI)1097-0215(19990820)84:4<421:AID-IJC16>3.0.CO;2-9.

4 Nakamura H, Kawasaki N, Taguchi M, Kabasawa K. Association of HER-2 overexpression with prognosis in nonsmall cell lung carcinoma: a metaanalysis. Cancer. 2005;103(9):1865–1873. doi:10.1002/cncr.20957.

5 Verri E, Guglielmini P, Puntoni M, et al. HER2/neu oncoprotein overexpression in epithelial ovarian cancer: evaluation of its prevalence and prognostic significance. Clinical study. Oncology. 2005;68(2-3):154–161. doi:10.1159/000086958.

6 Pegram M, Slamon D. Biological rationale for HER2/neu (c-erbB2) as a target for monoclonal antibody therapy. Semin Oncol. 2000;27(5 Suppl 9):13–19.

7 Serrano-Olvera A, Dueñas-González A, Gallardo-Rincón D, Candelaria M, La Garza-Salazar J de. Prognostic, predictive and therapeutic implications of HER2 in invasive epithelial ovarian cancer. Cancer Treatment Reviews. 2006;32(3):180–190. doi:10.1016/j.ctrv.2006.01.001.

8 Chen J-S, Lan K, Hung M-C. Strategies to target HER2/neu overexpression for cancer therapy. Drug Resist Updat. 2003;6(3):129–136. doi:10.1016/s1368-7646(03)00040-2.

9 Slamon DJ, Leyland-Jones B, Shak S, et al. Use of chemotherapy plus a monoclonal antibody against HER2 for metastatic breast cancer that overexpresses HER2. N Engl J Med. 2001;344(11):783–792. doi:10.1056/NEJM200103153441101.

10 Casi G, Neri D. Antibody-drug conjugates: basic concepts, examples and future perspectives. J Control Release. 2012;161(2):422–428. doi:10.1016/j.jconrel.2012.01.026.

11 Shastry M, Gupta A, Chandarlapaty S, Young M, Powles T, Hamilton E. Rise of Antibody-Drug Conjugates: The Present and Future. Am Soc Clin Oncol Educ Book. 2023;43:e390094. doi:10.1200/EDBK_390094.

12 Grairi M, Le Borgne M. Antibody-drug conjugates: prospects for the next generation. Drug Discov Today. 2024;29(12):104241. doi:10.1016/j.drudis.2024.104241.

13 Mckertish CM, Kayser V. Advances and Limitations of Antibody Drug Conjugates for Cancer. Biomedicines. 2021;9(8). doi:10.3390/biomedicines9080872.

14 Bagshawe KD. Antibody-directed enzyme prodrug therapy (ADEPT) for cancer. Expert Rev Anticancer Ther. 2006;6(10):1421–1431. doi:10.1586/14737140.6.10.1421.

15 Thuenemann EC, L. DHT, Lomonossoff GP, Steinmetz NF. Bluetongue Virus Particles as Nanoreactors for Enzyme Delivery and Cancer Therapy. Mol Pharm. 2021;18(3):1150–1156. doi:10.1021/acs.molpharmaceut.0c01053.

16 Qin M, Du G, Sun X. Biomimetic cell-derived nanocarriers for modulating immune responses. Biomater Sci. 2020;8(2):530–543. doi:10.1039/c9bm01444f.

17 Sutter M, Boehringer D, Gutmann S, et al. Structural basis of enzyme encapsulation into a bacterial nanocompartment. Nat Struct Mol Biol. 2008;15(9):939–947. doi:10.1038/nsmb.1473.

18 Giessen TW. Encapsulins: microbial nanocompartments with applications in biomedicine, nanobiotechnology and materials science. Curr Opin Chem Biol. 2016;34:1–10. doi:10.1016/j.cbpa.2016.05.013.

19 Nichols RJ, Cassidy-Amstutz C, Chaijarasphong T, Savage DF. Encapsulins: molecular biology of the shell. Crit Rev Biochem Mol Biol. 2017;52(5):583–594. doi:10.1080/10409238.2017.1337709.

20 Moon H, Lee J, Min J, Kang S. Developing genetically engineered encapsulin protein cage nanoparticles as a targeted delivery nanoplatform. Biomacromolecules. 2014;15(10):3794–3801. doi:10.1021/bm501066m.

21 Tamura A, Fukutani Y, Takami T, et al. Packaging guest proteins into the encapsulin nanocompartment from Rhodococcus erythropolis N771. Biotechnol Bioeng. 2015;112(1):13–20. doi:10.1002/bit.25322.

22 Giessen TW, Silver PA. Widespread distribution of encapsulin nanocompartments reveals functional diversity. Nat Microbiol. 2017;2:17029. doi:10.1038/nmicrobiol.2017.29.

23 Gabashvili AN, Chmelyuk NS, Efremova MV, Malinovskaya JA, Semkina AS, Abakumov MA. Encapsulins-Bacterial Protein Nanocompartments: Structure, Properties, and Application. Biomolecules. 2020;10(6). doi:10.3390/biom10060966.

24 van de Steen A, Wilkinson HC, Dalby PA, Frank S. Encapsulation of Transketolase into In Vitro-Assembled Protein Nanocompartments Improves Thermal Stability. ACS Appl Bio Mater. 2024;7(6):3660–3674. doi:10.1021/acsabm.3c01153.

25 Giessen TW. Encapsulins. Annu Rev Biochem. 2022;91:353–380. doi:10.1146/annurev-biochem-040320-102858.

26 Rennie C, Sives C, Boyton I, et al. In Vivo Behavior of Systemically Administered Encapsulin Protein Nanocages and Implications for their use in Targeted Drug Delivery. Advanced Therapeutics. 2024;7(2). doi:10.1002/adtp.202300360.

27 Mendes M, Mahita J, Blazeska N, et al. IEDB-3D 2.0: Structural data analysis within the Immune Epitope Database. Protein Sci. 2023;32(4):e4605. doi:10.1002/pro.4605.

28 Williams EM, Jung SM, Coffman JL, Lutz S. Pore Engineering for Enhanced Mass Transport in Encapsulin Nanocompartments. ACS Synth Biol. 2018;7(11):2514–2517. doi:10.1021/acssynbio.8b00295.

29 Adamson LSR, Tasneem N, Andreas MP, et al. Pore structure controls stability and molecular flux in engineered protein cages. Sci Adv. 2022;8(5):eabl7346. doi:10.1126/sciadv.abl7346.

30 Bae Y, Kim GJ, Kim H, Park SG, Jung HS, Kang S. Engineering Tunable Dual Functional Protein Cage Nanoparticles Using Bacterial Superglue. Biomacromolecules. 2018;19(7):2896–2904. doi:10.1021/acs.biomac.8b00457.

31 Michel-Souzy S, Hamelmann NM, Zarzuela-Pura S, Paulusse JMJ, Cornelissen JJLM. Introduction of Surface Loops as a Tool for Encapsulin Functionalization. Biomacromolecules. 2021;22(12):5234–5242. doi:10.1021/acs.biomac.1c01156.

32 Moon H, Lee J, Kim H, Heo S, Min J, Kang S. Genetically engineering encapsulin protein cage nanoparticle as a SCC-7 cell targeting optical nanoprobe. Biomater Res. 2014;18:21. doi:10.1186/2055-7124-18-21.

33 Lagoutte P, Mignon C, Stadthagen G, et al. Simultaneous surface display and cargo loading of encapsulin nanocompartments and their use for rational vaccine design. Vaccine. 2018;36(25):3622–3628. doi:10.1016/j.vaccine.2018.05.034.

34 Nygren P-A. Alternative binding proteins: affibody binding proteins developed from a small three-helix bundle scaffold. FEBS J. 2008;275(11):2668–2676. doi:10.1111/j.1742-4658.2008.06438.x.

35 Altai M, Liu H, Ding H, et al. Affibody-derived drug conjugates: Potent cytotoxic molecules for treatment of HER2 over-expressing tumors. J Control Release. 2018;288:84–95. doi:10.1016/j.jconrel.2018.08.040.

36 Ding H, Xu T, Zhang J, et al. Affibody-Derived Drug Conjugates Targeting HER2: Effect of Drug Load on Cytotoxicity and Biodistribution. Pharmaceutics. 2021;13(3). doi:10.3390/pharmaceutics13030430.

37 Li S, Jin Y, Su Y, et al. Anti-HER2 Affibody-Conjugated Photosensitizer for Tumor Targeting Photodynamic Therapy. Mol Pharm. 2020;17(5):1546–1557. doi:10.1021/acs.molpharmaceut.9b01247.

38 Sörensen J, Velikyan I, Sandberg D, et al. Measuring HER2-Receptor Expression In Metastatic Breast Cancer Using 68GaABY-025 Affibody PET/CT. Theranostics. 2016;6(2):262–271. doi:10.7150/thno.13502.

39 Hu X, Li D, Fu Y, et al. Advances in the Application of Radionuclide-Labeled HER2 Affibody for the Diagnosis and Treatment of Ovarian Cancer. Front Oncol. 2022;12:917439. doi:10.3389/fonc.2022.917439.

40 Miao H, Sun Y, Jin Y, Hu X, Song S, Zhang J. Application of a Novel 68Ga-HER2 Affibody PET/CT Imaging in Breast Cancer Patients. Front Oncol. 2022;12:894767. doi:10.3389/fonc.2022.894767.

41 Lan K-H, Tsai C-L, Chen Y-Y, et al. Affibody-conjugated 5-fluorouracil prodrug system preferentially targets and inhibits HER2-expressing cancer cells. Biochem Biophys Res Commun. 2021;582:137–143. doi:10.1016/j.bbrc.2021.09.078.

42 La Rodríguez de Fuente L, Cancela IG, Estévez-Salguero ÁM, Iglesias P, Costoya JA. Development of a biosensor based on a new marine luciferase fused to an affibody to assess Her2 expression in living cells. Anal Chim Acta. 2022;1221:340084. doi:10.1016/j.aca.2022.340084.

43 Zhao Z, Tao X, Xie Y, et al. In Situ Prodrug Activation by an Affibody-Ruthenium Catalyst Hybrid for HER2-Targeted Chemotherapy. Angew Chem Int Ed Engl. 2022;61(26):e202202855. doi:10.1002/anie.202202855.

44 Smith B, Lyakhov I, Loomis K, et al. Hyperthermia-triggered intracellular delivery of anticancer agent to HER2(+) cells by HER2-specific affibody (ZHER2-GS-Cys)-conjugated thermosensitive liposomes (HER2(+) affisomes). J Control Release. 2011;153(2):187–194. doi:10.1016/j.jconrel.2011.04.005.

45 Shipunova VO, Sogomonyan AS, Zelepukin IV, Nikitin MP, Deyev SM. PLGA Nanoparticles Decorated with Anti-HER2 Affibody for Targeted Delivery and Photoinduced Cell Death. Molecules. 2021;26(13). doi:10.3390/molecules26133955.

46 Zhang Y, Jiang S, Zhang D, Bai X, Hecht SM, Chen S. DNA-affibody nanoparticles for inhibiting breast cancer cells overexpressing HER2. Chem Commun (Camb). 2017;53(3):573–576. doi:10.1039/c6cc08495h.

47 Zmyslia M, Capper MJ, Grimmeisen M, et al. A nanoengineered tandem nitroreductase: designing a robust prodrug-activating nanoreactor. RSC Chem Biol. 2024. doi:10.1039/d4cb00127c.

48 Altenburg WJ, Rollins N, Silver PA, Giessen TW. Exploring targeting peptide-shell interactions in encapsulin nanocompartments. Sci Rep. 2021;11(1):4951. doi:10.1038/s41598-021-84329-z.

49 Lohner P, Zmyslia M, Thurn J, et al. Inside a Shell—Organometallic Catalysis Inside Encapsulin Nanoreactors. Angewandte Chemie. 2021;133(44):24028–24034. doi:10.1002/ange.202110327.

50 Putri RM, Allende-Ballestero C, Luque D, et al. Structural Characterization of Native and Modified Encapsulins as Nanoplatforms for in Vitro Catalysis and Cellular Uptake. ACS Nano. 2017;11(12):12796–12804. doi:10.1021/acsnano.7b07669.

51 Wikman M, Steffen A-C, Gunneriusson E, et al. Selection and characterization of HER2/neu-binding affibody ligands. Protein Eng Des Sel. 2004;17(5):455–462. doi:10.1093/protein/gzh053.

52 Orlova A, Magnusson M, Eriksson TLJ, et al. Tumor imaging using a picomolar affinity HER2 binding affibody molecule. Cancer Res. 2006;66(8):4339–4348. doi:10.1158/0008-5472.CAN-05-3521.

53 Liu H, Seijsing J, Frejd FY, Tolmachev V, Grôslund T. Target-specific cytotoxic effects on HER2-expressing cells by the tripartite fusion toxin ZHER2:2891-ABD-PE38X8, including a targeting affibody molecule and a half-life extension domain. International Journal of Oncology. 2015;47(2):601–609. doi:10.3892/ijo.2015.3027.

54 Dong D, Xia G, Li Z, Li Z. Human Serum Albumin and HER2-Binding Affibody Fusion Proteins for Targeted Delivery of Fatty Acid-Modified Molecules and Therapy. Mol Pharm. 2016;13(10):3370–3380. doi:10.1021/acs.molpharmaceut.6b00265.

55 ERBB2 DepMap Gene Summary. https://depmap.org/portal/gene/ERBB2?tab=characterization&characterization=expression. Updated January 28, 2025. Accessed January 28, 2025.

56 Schmitt JF, Susil BJ, Hearn MT. Aberrant FGF-2, FGF-3, FGF-4 and C-erb-B2 gene copy number in human ovarian, breast and endometrial tumours. Growth Factors. 1996;13(1-2):19–35. doi:10.3109/08977199609034564.

57 Giessen TW. The Structural Diversity of Encapsulin Protein Shells. Chembiochem. 2024;25(24):e202400535. doi:10.1002/cbic.202400535.

58 Ross J, McIver Z, Lambert T, et al. Pore dynamics and asymmetric cargo loading in an encapsulin nanocompartment. Sci Adv. 2022;8(4):eabj4461. doi:10.1126/sciadv.abj4461.

59 Kwon S, Andreas MP, Giessen TW. Pore Engineering as a General Strategy to Improve Protein-Based Enzyme Nanoreactor Performance. ACS Nano. 2024;18(37):25740–25753. doi:10.1021/acsnano.4c08186.

