## Supporting Information for "Customized Protein Nanoreactors for Affibody-Directed Activation of 5-Fluorcytosin in HER2-Positive Cells"

### General materials and methods

#### Chemicals, analytics and general remarks

Reagents were purchased from commercial suppliers (Sigma-Aldrich, Carl Roth, BLDpharm, Thermo Fischer Scientific, New England Biolabs, Promega) and used as received, unless noted otherwise. Ampicillin, lysozyme, dithiothreitol (DTT), phenylmethylsulfonyl fluoride (PMSF), phenazine-methosulfate (PMS), dimethyl sulfoxide (DMSO) were acquired from Sigma-Aldrich, Germany. Acetonitrile (ACN), HEPES, Tris-HCl, sodium chloride (NaCl), imidazole, Tween 20 and ethylenediaminetetraacetic acid (EDTA) were purchased from Carl Roth, Germany. 5-Fluorocytosine and desthiobiotin were purchased from BLDpharm. CellMask Orange plasma membrane stain and 5-fluorouracile were obtained from Thermo Fischer Scientific. CellTiter 96 AQueous MTS Reagent Powder was purchased from Promega, Germany. HPLC chemicals and solvents were obtained in analytical grade and used as received.

The pETDuet-1 and pCDFDuet-1 plasmid used for cloning were obtained from Novagen, Merck. Restriction enzymes were sourced from New England Biolabs, USA.

Encapsulin: UniProt ID: I7G8Y9, Organism: *Mycobacterium smegmatis* mc$^2$155, Other descriptors: 29 kDa antigen CFP29, MSMEI_5672

ELS amino acids sequence: Ser-Leu-Gly-Ile-Gly-Ser-Leu-Lys-Gly-Thr-Arg.

#### Bacterial strains and cell lines

| | | |
|---|---|---|
| **Bacterial strains** | | |
| *E. coli* XL1-Blue | Host strain for cloning | Stratagene/Agilent |
| *E. coli* BL21 Star (DE3) | Host strain for protein expression | Novagen/Merck |
| **Mammalian cell lines** | | |
| J774A.1 | Murine (monocyte-) macrophage cell line | BIOSS, University of Freiburg |
| SKOV-3 | Human ovarian cancer cell line (HER2$^+$) | Gift from Prof. Tilman Brummer, University of Freiburg |
| OV7 | Human ovarian cancer cell line (HER2$^-$) | Gift from Dr. Elisabetta Grillo, University of Brescia |

### Genetic engineering

#### Cloning of pETDuet-1: Enc-Strep-Tag (MCS1); tdCD-ELS (MCS2)

The commercially available pETDuet-1 vector (Novagen) was modified to include a Strep-tag II. This was achieved by introducing the tag sequence into MCS1 via an annealed oligonucleotide cloning method, utilizing the primer pair ZM1_fw/ZM2_rev (as indicated in Table S1), which was inserted between the SacI and HindIII sites. The open reading frame encoding MSMEI_5672 was subsequently amplified from the genomic DNA of *M. smegmatis* mc$^2$155 using the primer pair ZM3_fw/ZM4_rev and then inserted into MCS1 between the NcoI and SacI restriction sites. Concurrently, the MCS2 was modified to incorporate the tdCD-ELS sequence, which was commercially synthesized. Standard cloning techniques were applied to insert this sequence into MCS2 using the primer pair ZM5_fw/ZM6_rev, along with NdeI and PacI restriction enzymes.

#### Cloning of pETDuet-1: Enc-Strep-Tag (MCS1); eGFP-ELS (MCS2)

For this construct the Enc-Strep-Tag was introduced into MCS1 following the same methodology as outlined previously. Subsequently, MCS2 was engineered to include the eGFP-ELS sequence, which was amplified from a commercially synthesized vector and inserted between the NdeI and PacI restriction sites.

**Cloning of pCDFDuet-1: Enc-ZHER$_{2:342}$-His-Tag (MCS1); Enc (MCS2)**

The commercially available pCDFDuet-1 vector (Novagen) underwent modifications. The Enc-ZHER$_{2:342}$[1]-His-Tag sequence was amplified from a commercially synthesized vector and inserted between the NcoI and AflII sites of MCS1. MCS2 was modified by inserting the open reading frame encoding MSMEI_5672, amplified using primer pair ZM7_fw/ZM8_rev, into MCS2 between the NdeI and PacI restriction sites.

**Cloning of pETDuet-1: tdCD-ELS (MCS2)**

MCS2 of the commercially available pETDuet-1 vector (Novagen) was modified to include the tdCD-ELS sequence. The sequence was amplified from the previously constructed pETDuet-1: Enc-Strep-Tag (MCS1); tdCD (MCS2) using primer pair ZM9_fw/ZM10_rev, and inserted using NdeI and PacI restriction enzymes.

**Cloning of pETDuet-1: eGFP-ELS (MCS2)**

Similarly, MCS2 of the commercially available pETDuet-1 vector (Novagen) was modified to add the eGFP-ELS sequence. This sequence was amplified from the pETDuet-1: Enc-Strep-Tag (MCS1); eGFP (MCS2) construct using primer pair ZM9_fw/ZM10_rev, and incorporated using NdeI and PacI restriction enzymes.

**Table S1. Primers and oligonucleotides used in this study.**

| Name | Sequence |
|---|---|
| ZM1_fw | CTGGAGCCACCCGCAGTTCGAAAAATGAA |
| ZM2_rev | AGCTTTCATTTTTCGAACTGCGGGTGGCTCCAGAGCT |
| ZM3_fw | TAAGCACCATGGGAAACAACCTCTATCGCGACCTCGC |
| ZM4_rev | TAAGCAGAGCTCGGGGGTCAGCGCGACAGAG |
| ZM5_fw | CTACATATGGTGACAGGGGGAATGGC |
| ZM6_rev | AGACTTAATTAATCAGCGGGTTCCTTTC |
| ZM7_fw | AGTCATATGGGAAACAACCTCTATCGCG |
| ZM8_rev | ATGTTAATTAATCAGGGGGTCAGCGCGAC |
| ZM9_fw | TTGTACACGGCCGCATAATC |
| ZM10_rev | GGTGGCAGCAGCCTAG |

**Complete protein sequences**

Purple: Strep-tag II; Orange: His-Tag; Blue: protein sequence encapsulin; Yellow: protein sequence CD; Green: protein sequence eGFP; Teal blue: protein sequence ZHER$_{2:342}$ affibody; Brown: linker between monomers; Red: encapsulin localization sequence.

Enc-Strep-Tag

MGNNLYRDLAPITESAWAEIELEATRTFKRHIAGRRVVDVSGPNGPTTASVSTGHLLDVSPPGDGVIAHLR
DAKPLVRLRVPFTVARRDIDDVERGSQDSDWDPVKDAAKKLAFVEDRAIFEGYAAASIEGIRSSSSNPALA
LPDDAREIPDVIAQALSELRLAGVDGPYSVLLSAETYTKVSETTAHGYPIREHINRLVDGEIIWAPAIDGAFV
LSTRGGDFDLQLGTDVSIGYLSHDAEVVHLYMEETMTFLCYTAEASVALTPELWSHPQFEK

tdCD

MVTGGMASKWDQKGMDIAYEEAALGYKEGGVPIGGCLINNKDGSVLGRGHNMRFQKGSATLHGEISTLE
NCGRLEGKVYKDTTLYTTLSPCDMCTGAIIMYGIPRCVVGENVNFKSKGEKYLQTRGHEVVVVDDERCKK
IMKQFIDERPQDWFEDIGELEGSAGQGAQAGQGAQAGSSAGTVTGGMASKWDQKGMDIAYEEAALGYK
EGGVPIGGCLINNKDGSVLGRGHNMRFQKGSATLHGEISTLENCGRLEGKVYKDTTLYTTLSPCDMCTGA

IIMYGIPRCVVGENVNFKSKGEKYLQTRGHEVVVVDDERCKKIMKQFIDERPQDWFEDIGELAGGGSASL
GIGSLKGTR

eGFP

MVSKGEELFTGVVPILVELDGDVNGHKFSVSGEGEGDATYGKLTLKFICTTGKLPVPWPTLVTTLTYGVQ
CFSRYPDHMKQHDFFKSAMPEGYVQERTIFFKDDGNYKTRAEVKFEGDTLVNRIELKGIDFKEDGNILGH
KLEYNYNSHNVYIMADKQKNGIKVNFKIRHNIEDGSVQLADHYQQNTPIGDGPVLLPDNHYLSTQSALSKD
PNEKRDHMVLLEFVTAAGITLGMDELYKAAAGSLGIGSLKGTR

Enc-ZHER$_{2:342}$-His-Tag

MGNNLYRDLAPITESAWAEIELEATRTFKRHIAGRRVVDVSGPNGPTTASVSTGHLLDVSPPGDGVIAHLR
DAKPLVRLRVPFTVARRDIDDVERGSQDSDWDPVKDAAKKLAFVEDRAIFEGYAAASIEGIRSSSSNPALA
LPDDAREIPDVIAQALSELRLAGVDGPYSVLLSAETYTKVSETTAHGYPIREHINRLVDGEIIWAPAIDGAFV
LSTRGGDFDLQLGTDVSIGYLSHDAEVVHLYMEETMTFLCYTAEASVALTPGTGGGGSGGGGSKLVDNK
FNKEMRNAYWEIALLPNLNNQQKRAFIRSLYDDPSQSANLLAEAKKLNDAQAPKAAALEHHHHHH

Enc

MGNNLYRDLAPITESAWAEIELEATRTFKRHIAGRRVVDVSGPNGPTTASVSTGHLLDVSPPGDGVIAHLR
DAKPLVRLRVPFTVARRDIDDVERGSQDSDWDPVKDAAKKLAFVEDRAIFEGYAAASIEGIRSSSSNPALA
LPDDAREIPDVIAQALSELRLAGVDGPYSVLLSAETYTKVSETTAHGYPIREHINRLVDGEIIWAPAIDGAFV
LSTRGGDFDLQLGTDVSIGYLSHDAEVVHLYMEETMTFLCYTAEASVALTP

**Protein production and purification**

Plasmids encoding the proteins of interest were transformed into chemically competent *E. coli* BL21 Star (DE3) cells. Transformant colonies were selected on LB-agar plates containing the respective antibiotics: ampicillin for pETDuet-1 constructs and streptomycin for pCDFDuet-1 constructs. A single colony was subsequently inoculated into LB medium supplemented with the corresponding antibiotic and cultured overnight at 37 °C with shaking at 180 rpm. Subsequently, a 5 mL pre-culture was inoculated into 1 L of auto-induction medium (also supplemented with the appropriate antibiotic) and incubated at 19 °C for 65 hours.

Cells were collected by centrifugation at 5000 × g for 40 minutes at 4 °C and resuspended in lysis buffer. For Strep-tagged proteins, the lysis buffer consisted of 100 mM Tris-HCl, 150 mM NaCl, 1 mM EDTA, pH 8, while for His-tagged proteins, it comprised 20 mM sodium phosphate, 500 mM NaCl, and 20 mM imidazole, pH 7.4. The lysis buffer was freshly supplemented with 1 mM DTT, 1 mM PMSF, and 1 mg/mL lysozyme, and cells were incubated on ice for 30 to 45 minutes. Cell lysis was achieved via sonication on ice using a Bandelin Sonopuls HD 2070 ultrasonic homogenizer (Bandelin Electronics, Berlin, Germany) for two cycles of 3 minutes on/off at 40% amplitude. Cell debris was removed by centrifugation at 15,000 × g for 30 minutes.

Proteins carrying a Strep-tag were purified using a 5 mL StrepTrap HP column (Cytiva, Germany) and eluted with an elution buffer comprising of 100 mM Tris-HCl, 150 mM NaCl, 1 mM EDTA, and 2.5 mM desthiobiotin, pH 8. A typical purification of Strep-tagged encapsulin from 1 L of culture yielded approximately 25 to 30 mg of protein after pooling all fractions from the 5 mL StrepTrap HP column (4.5 to 5 mg/mL).

Proteins bearing a His-tag were purified using a 5 mL His-Trap HP column. Following initial washing, proteins were eluted using a stepwise gradient of elution buffer (20 mM sodium phosphate, 500 mM NaCl, 500 mM imidazole, pH 7.4) at concentrations of 20%, 60%, and 100%. Only the fractions eluted with 100% elution buffer were retained for subsequent applications. A typical purification of His-tagged encapsulin from 1 L of culture yielded approximately 10-15 mg of protein, with all fractions eluted at 100% elution buffer from the 5 mL His-Trap HP column pooled (3–3.5 mg/mL).

Further purification was achieved using size exclusion chromatography on a HiLoad Superdex 200 PG column (Äkta Protein Purification System, GE Healthcare, Germany). Proteins were eluted from the column with an elution buffer of 50 mM HEPES, pH 7.4. Fractions were analyzed by SDS-PAGE, pooled, subdivided, and stored at -80 °C for future use.

**Biophysical characterization**

**Negative stain-transmission electron microscopy (TEM)**

Nanocompartment samples were diluted to 0.2 mg/mL in HEPES buffer (50 mM, pH 7.4) and applied for 1 minute onto carbon-coated grids (Electron Microscopy Sciences, FCF300-CU), which had been previously glow-discharged for 25 s. Following adsorption, grid was rinsed with water and subsequently stained with 1% (w/v) uranyl acetate for 20 s. Excess liquid was removed using filter paper. Visualization was carried out on a Thermo Scientific Talos L120C transmission electron microscope, operated at 120 kV. Image processing and particle size analysis were performed in ImageJ.

**Dynamic light scattering (DLS)**

DLS analysis was conducted at 25°C on a Zetasizer Nano ZS (Malvern Instruments, Malvern, UK) using ultra-low volume quartz cuvette (ZEN2112) containing 1 mg/mL of nanocompartment sample in 50 mM HEPES, 0.05% Tween 20, pH 7.4. Each specimen was measured in three consecutive runs under the following conditions: attenuator: 8; mean count rate (kcps): 260 - 420. The data were analyzed and interpreted employing the Zetasizer Nanoseries software (Malvern Instruments, Malvern, UK) with the general-purpose model. For each measurement, the intensity-weighted mean hydrodynamic diameter was computed and documented.

**Asymmetric flow field flow fractionation (AF4)**

AF4 system consisted of a flow controller (Eclipse AF4, Wyatt, Dernbach, Germany), a MALS detector (DAWN Heleos II, Wyatt), a differential refractometer (Optilab T-rEX, Wyatt) and the separation channel (SC channel, regenerated cellulose membrane, cut-off 10 kDa, 350 µm spacer, wide type, Wyatt). The elution buffer used was 50 mM HEPES adjusted to pH 7.4 and filtered through 0.1 µm. AF4 runs were conducted at 22 °C with a channel flow rate of 1.0 mL/min and the following flow sequence (Vx = cross flow in mL/min): (a) elution (2 min, Vx: 1.0); (b) focus (1 min, Vx: 1.5), focus + inject (1 min, Vx: 1.5, inject flow: 0.2 mL/min); focus (2 min, Vx: 1.5); (c) elution (30 min, Vx: 1.0); elution (30 min, linear Vx gradient: 1.0 to 0.05); (d) elution (5 min, Vx: 0.0); (e) elution + inject (2 min, Vx: 0.0). A total protein mass of 23 ± 2.0 µg was injected. The eluted sample concentration was calculated from the refractive index signal using a specific refractive index increment of 0.185 mL/g. Mass-weighted mean values of the molar masses and the radii of gyration were calculated from MALS data using the ASTRA 8 software package (Wyatt).

**Blue Native PAGE**

Native protein nanocompartments were analyzed on NativePAGE 3–12% Bis-Tris 1.0 mm Mini Protein Gels (Thermo Fisher) alongside the NativeMark™ Unstained Protein Standard (Thermo Fisher). Protein samples contained 2.5 µg of protein in NativePAGE Sample Buffer. Electrophoresis buffers (anode, dark cathode, and light cathode) were prepared according to the manufacturer's protocol (Thermo Fisher - NativePAGE Bis-Tris Gel Manual). Separation was initiated in dark blue cathode buffer at 120 V for 25 min, after which the buffer was exchanged to light blue cathode buffer and the gel run for approximately 12 h at 50 V. Voltage was then sequentially increased to 120 V for 1 h and to 200 V for a final 30 min. Gels were stained in Coomassie Brilliant Blue G solution (80 mg/L, 30 mM HCl) and destained in water.

**SDS-PAGE and gel densitometry**

Protein samples (20 µL) were combined with an equal volume (20 µL) of 2 × SDS sample buffer (125 mM Tris–HCl (pH 6.8), 4% SDS, 20% (v/v) glycerol, 0.02% bromophenol blue, 200 mM DTT) and then incubated at 95 °C for 10 minutes for complete protein denaturation. Polyacrylamide gels (15%) were cast in the laboratory and loaded into the vertical gel electrophoresis system. Prior to sample application, both upper and lower reservoirs were filled with SDS-PAGE running buffer (25 mM Tris, 186 mM Glycine, 0.1% SDS). An equal volume for each sample was applied into the wells. The BlueStar Plus Prestained Protein Marker (MWP04, Nippon), covering a molecular weight range of 10–240 kDa, served as the standard for monitoring separation. Electrophoresis proceeded at 200 V and 60 mA for approximately 1 hour, until bromophenol blue reached the bottom of the separating gel.

Post-run, gels were submerged overnight in staining solution containing 0.1% Coomassie R-250 in 10% (w/v) ammonium sulfate, 1% (v/v) phosphoric acid, and 10% (v/v) methanol. Destaining was performed with distilled water. Gel images were acquired using the UVP ChemStudio Touch 815 (Analytik Jena).

Densitometric analysis was executed with ImageJ software. Background signal was subtracted for each lane, and regions of interest (ROIs) were defined for bands corresponding to encapsulin monomer (Enc, 28.8 kDa) and affibody-fused encapsulin monomer ($^{ZHER}$Enc, 37.8 kDa), particularly for $^{ZHER}$Enc{tdCD} and $^{ZHER}$Enc{eGFP} samples. Band intensities were quantified and presented graphically. To accurately quantify the ratio between Enc and $^{ZHER}$Enc, a correction factor was applied to account for the difference in molecular weights, as larger proteins bind more Coomassie dye and therefore produce a stronger signal. Specifically, the signal intensity for the Enc band was multiplied by a factor of 1.3125 (calculated as 37.8 kDa / 28.8 kDa). This adjustment allowed for accurate comparison of the relative abundance of the two subunit types. Based on the corrected signal ratio, the number of Enc and $^{ZHER}$Enc subunits per shell was calculated, considering that each encapsulin nanocompartment is composed of exactly 60 monomers. The experiment was performed in duplicate to ensure reproducibility.

**Cellular analysis**

**Cell lines and culture conditions**

The mouse monocyte cell line J774A.1 and human ovarian cancer cell lines SKOV-3 and OV7 were maintained under standard culture conditions at 37 °C and 5% CO2. SKOV-3 and J774A.1 cells were cultured in high-glucose DMEM supplemented with 10% fetal calf serum and 10% heat-inactivated fetal calf serum, respectively. The OV7 cell line was cultured in a DMEM:HAMS F-12 medium supplemented with 5% fetal calf serum, 0.5 µg/mL hydrocortisone, and 10 µg/mL insulin.

**Fluorescence microscopy**

For live cell imaging, a 300 µL suspension of 1 × $10^5$ SKOV-3 cells was seeded into each well of µ-Slide 8-well chambers with polymer coverslip bottoms (Ibidi, Germany) and allowed to attach overnight. After the attachment period, the medium was replaced with 300 µL of fresh medium containing either 20 nM Enc{eGFP}, 20 nM $^{ZHER}$Enc{eGFP}, or was left untreated. The cells were then incubated for 90 minutes. Following the incubation, the medium was removed, and the cells were washed twice with serum-free DMEM. The plasma membrane was subsequently stained with CellMask Orange (2.5 µg/mL) prior to imaging (red fluorescence). Fluorescence imaging was conducted using an inverted Zeiss LSM 880 laser scanning microscope equipped with an LD LCI Plan-Apochromat 63×/1.40 Oil DIC M27 objective. Green fluorescence (eGFP) was detected between 499-552 nm upon excitation at 488 nm, while red fluorescence (CellMask Orange) was excited at 561 nm and emission was collected between 566-735 nm. All images were captured under consistent settings to ensure comparability across different experimental conditions.

**Flow cytometry**

SKOV-3 and OV7 cells were cultivated in T75 flasks and subsequently harvested for flow cytometry. Initially, each flask was rinsed twice with 6 mL of PBS, followed by the addition of 3 mL of Trypsin-EDTA solution and incubation at 37 °C for 5 minutes to facilitate cell detachment. Next, 7 mL of cell culture medium was added, and the resulting cell suspension was transferred to 15 mL Falcon tubes. The cells were pelleted by centrifugation at 200 × g at room temperature, washed twice with warm PBS, and counted. The final cell pellets were resuspended in ice-cold FACS buffer (2% v/v FBS and 0.08% w/v sodium azide in PBS).

Aliquots of $1 \times 10^6$ cells in 70 µL of FACS buffer were incubated on ice for 60 minutes with 30 µL of protein solution (final volume: 100 µL). Cells were treated with either Enc{eGFP} or $^{ZHER}$Enc{eGFP} at three different amounts (3.3 pmol, 10 pmol, and 20 pmol), or left untreated as a control. Protein dilutions were prepared in 50 mM HEPES (pH 7.4) to achieve the desired concentrations. Following incubation, the cells were washed twice with 1 mL of FACS buffer, resuspended in 400 µL of FACS buffer, and stained with 4′,6-diamidin-2-phenylindole (DAPI, final concentration: 0.15 µg/mL) for 5 minutes to exclude non-viable cells. All samples were analyzed within one hour on a BD LSRFortessa flow cytometer (BD, Germany) with 50 000 events being recorded per sample. Data processing was performed using FlowJo v10 software. Single cells were gated by plotting FSC-W versus FSC-A to eliminate cell aggregates, and DAPI-positive events were excluded to isolate the live-cell population. This final population was then used for subsequent analysis of protein binding.

**Functional assays**

**In vitro prodrug activation**

In vitro prodrug activation was conducted in HEPES buffer (50 mM, pH 7.4) at 37 °C and 350 rpm. The reaction was initiated by adding a dilution of Enc{tdCD} enzyme to achieve a final concentration of 188 µg/mL (corresponds to 75 nM) or by adding assay buffer for the negative control, both mixed with 5-FC dilution at a final concentration of 3 mM. After 1 hour of incubation, aliquots from both control and reaction samples were collected, and the reactions were terminated by mixing 1:1 with cold acetonitrile (ACN).

Chromatographic separation was performed using an Agilent 1290 Infinity II UHPLC system (Agilent, Germany) equipped with an ACQUITY UPLC BEH Amide column (130 Å, 1.7 µm, 150 x 2.1 mm, Waters, USA). The mobile phases included A: water, B: 0.5% ammonium hydroxide in water, and C: ACN. The analysis was conducted at a flow rate of 0.5 mL/min and a column temperature of 40 °C, following a gradient program that began with 0% A, 5% B, and 95% C, transitioning to 45% A, 5% B, and 50% C, over the first 6 minutes. The system was maintained at 45% A, 5% B, and 50% C from 6 to 7 minutes. Mass spectrometric analysis was performed using an Agilent 6545 LC/Q-TOF system operating in negative ion mode.

**In cellulo prodrug activation**

To investigate prodrug activation in the J774A.1 cell line, 2000 cells/well were plated in 96-well plates. After allowing the cells to attach overnight, the medium was replaced with either 20 nM Enc{eGFP} or 20 nM Enc{tdCD} protein dilutions in complete culture medium, or with fresh culture medium for the untreated control group. After 18 hours of incubation, the cells were washed twice with 200 µL PBS, and freshly prepared 0.5 mM 5-FC in complete culture medium (filter-sterilized) was added. At identical intervention time points, the control wells received complete culture medium instead of protein dilutions, were washed twice with PBS, and had either 0.5 mM 5-FC (negative control) or 0.5 mM 5-FU (positive control) added. After 72 hours of incubation, cell viability was assessed using the CellTiter 96® AQueous Cell Proliferation Assay (MTS assay, Promega, Germany) according to the manufacturer's instructions. Briefly, CellTiter 96® AQueous MTS Reagent Powder was dissolved in DPBS at 1 mg/mL, filter-sterilized, aliquoted, and stored at -20 °C. Phenazine methosulfate (PMS; Sigma) was prepared at a concentration of 0.92 mg/mL in DPBS, also filter-sterilized through a 0.2 µm filter, aliquoted, and stored at -20 °C. Immediately before performing the MTS assay, 1.524 mL of the MTS solution was combined with 76 µL of the PMS solution. An additional 8 mL of medium was added to this mixture. In the assay plate, the old medium was replaced with 120 µL of the MTS/PMS solution mixture. Furthermore, 120 µL of the same MTS/PMS mixture in medium was added to empty wells as a negative control. The cells were incubated for 2.5 to 3 hours under standard culture conditions, after which the plate was briefly shaken and measured using a microplate reader (Tecan Spark® Multimode Microplate Reader) at an optical density of 490 nm. All conditions were assessed in triplicate, and the experiment was repeated twice. The average background absorbance (from “no cell” control wells) was subtracted from all values to obtain corrected absorbance values. The absorbance of untreated cells was set at 100% viability. The viability of experimental wells was calculated by dividing their absorbance by the absorbance of untreated cells and multiplying by 100%.

SKOV-3 cells were seeded in a 96-well plate at a density of 6000 cells/well and allowed to attach overnight. The following day, the medium was replaced with either 50 nM $^{ZHER}$Enc{eGFP} or 50 nM $^{ZHER}$Enc{tdCD} protein dilutions in complete culture medium and incubated for 1 hour. After incubation, the cells were washed twice with 200 µL PBS, and freshly prepared 4 mM 5-FC in complete culture medium (filter-sterilized) was added. At the same intervention time points, control wells received complete culture medium instead of protein dilutions, were washed twice with PBS, and then had either 4 mM 5-FC (negative control) or 2 mM 5-FU (positive control) added. Dilutions exceeding 2 mM of 5-FU could not be prepared in the medium due to solubility limitations. After 120 hours of incubation, cell viability was determined using the MTS assay as described above.

### Supporting figures

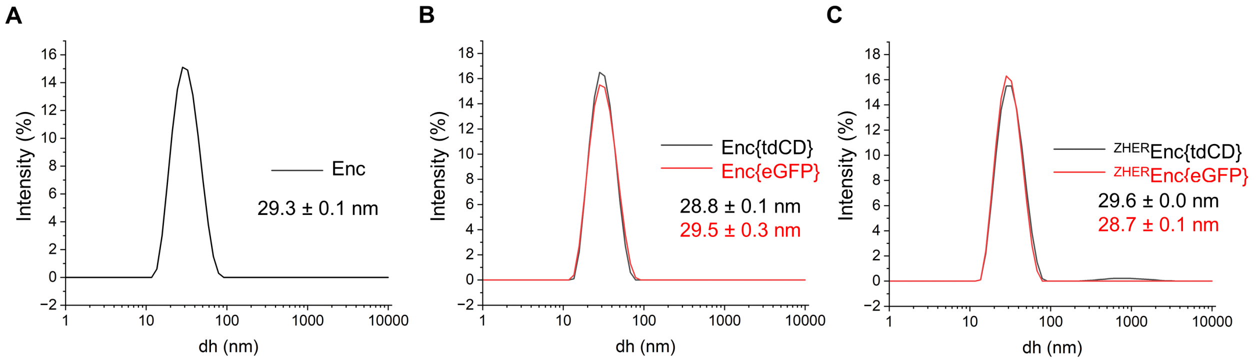

**Figure S1:** DLS analysis of encapsulin constructs: Enc (A), Enc{tdCD} and Enc{eGFP} (B), $^{ZHER}$Enc{tdCD} and $^{ZHER}$Enc{eGFP} (C).

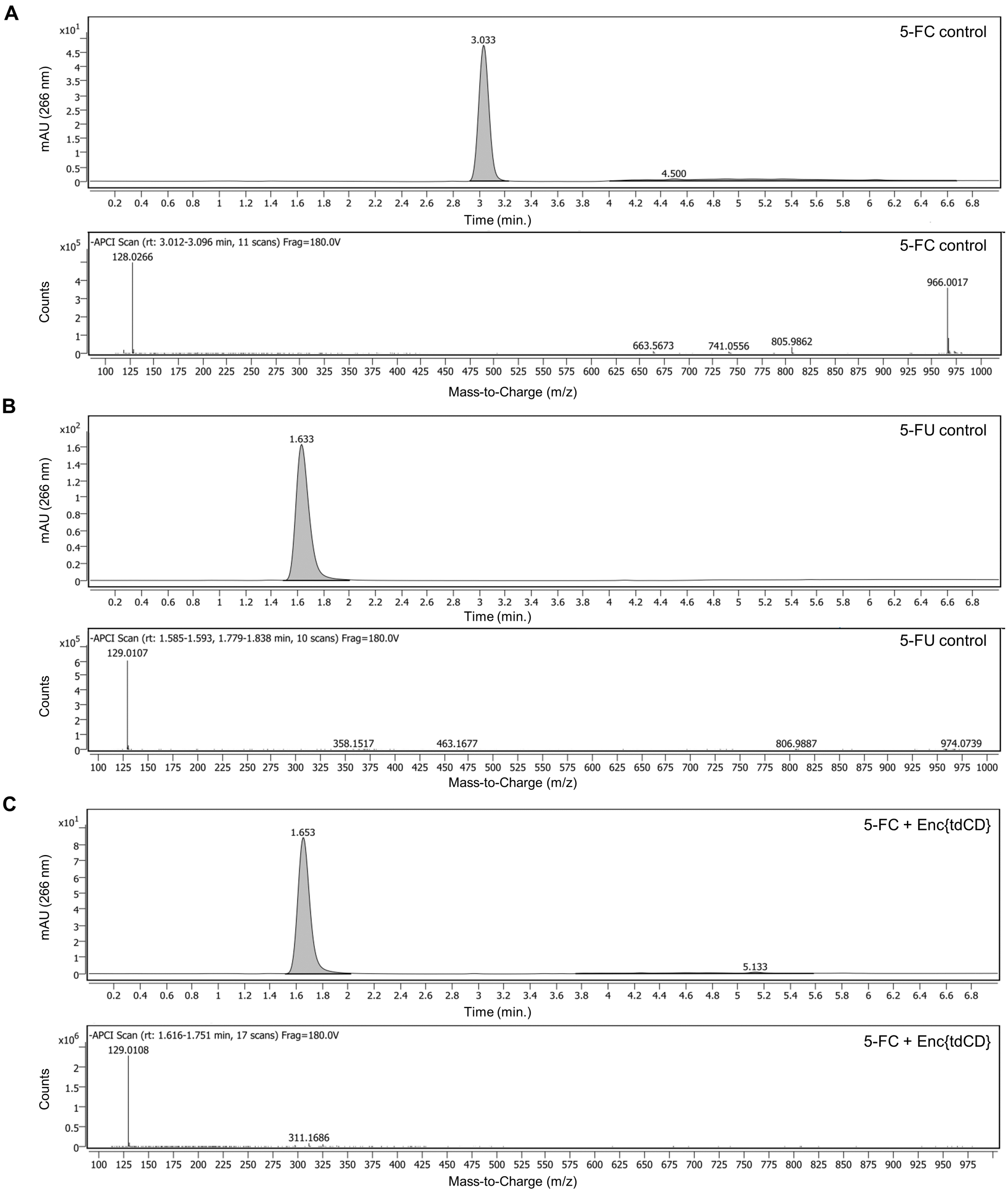

**Figure S2:** HPLC chromatograms and APCI-MS spectra of 5-FC (A), 5-FU (B), and Enc{tdCD}-mediated conversion of 5-FC to 5-FU (C). 5-FC expected mass = 128.03 Da (M – H$^+$); 5-FU expected mass = 129.02 Da (M – H$^+$).

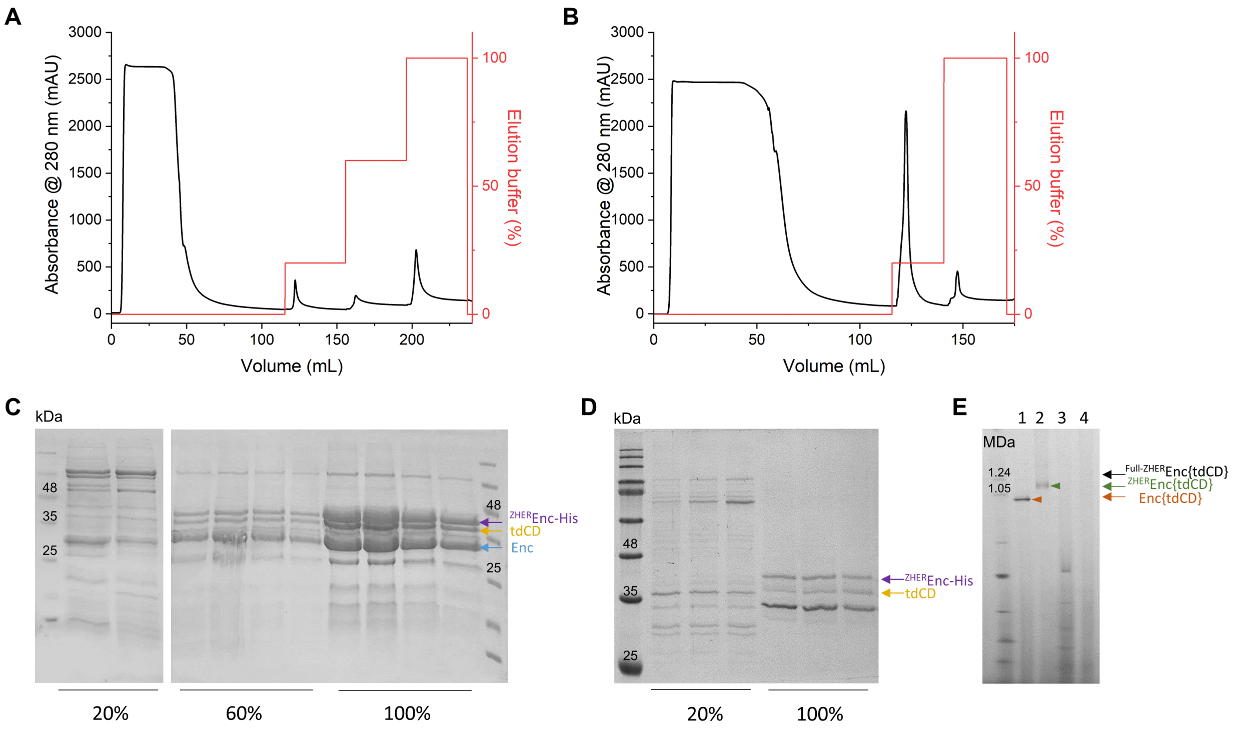

**Figure S3**: Purification and assembly analysis of $^{\text{ZHER}}$Enc{tdCD} and $^{\text{Full-ZHER}}$Enc{tdCD}. (A, B) Representative chromatograms of affinity chromatography runs on a HisTrap HP column for $^{\text{ZHER}}$Enc{tdCD} (A) and $^{\text{Full-ZHER}}$Enc{tdCD} (B). Stepwise elution was performed to enhance sample homogeneity by fractionating protein subpopulations according to their His-tag content, which directly correlates with affibody incorporation. For $^{\text{ZHER}}$Enc{tdCD}, whose shell consists of a mixed population of wild-type and affibody-fused encapsulin monomers, a three-step elution was applied (20%, 60%, and 100% elution buffer) to fractionate protein populations with varying His-tag density. In contrast, $^{\text{Full-ZHER}}$Enc{tdCD}, whose shell consists exclusively of affibody-fused encapsulin monomers, was purified using a two-step elution (20% and 100%), as no heterogeneity in shell composition was expected. (C, D) SDS-PAGE analysis of elution fractions from affinity chromatography for $^{\text{ZHER}}$Enc{tdCD} (C) and $^{\text{Full-ZHER}}$Enc{tdCD} (D). At 20% elution, non-target proteins and impurities are removed, while protein populations with a lower proportion of affibody-fused encapsulin monomers in the shell of $^{\text{ZHER}}$Enc{tdCD} elute at 60%. The remaining target protein, enriched in affibody-fused encapsulin monomers, elutes at 100%, ensuring higher sample homogeneity. The differential elution behavior of $^{\text{ZHER}}$Enc{tdCD} reflects the variability in affibody incorporation, as only a fraction of the shell protomers carries a His-tag. In contrast, the shell of $^{\text{Full-ZHER}}$Enc{tdCD} is composed entirely of His-tagged subunits, warranting the use of a two-step elution strategy. Protein bands are indicated as follows: affibody-fused encapsulin monomer (purple arrow, 37.8 kDa), wild-type encapsulin monomer (blue arrow, 28.8 kDa), and tdCD (yellow arrow, 38.4 kDa). (E) BN-PAGE analysis: lane 1 – Enc{tdCD}, lane 2 – $^{\text{ZHER}}$Enc{tdCD}, lane 3 – $^{\text{Full-ZHER}}$Enc{tdCD} 20% elution, lane 4 – $^{\text{Full-ZHER}}$Enc{tdCD} 100% elution. The absence of high-molecular-weight species in $^{\text{Full-ZHER}}$Enc{tdCD} (black arrows indicate expected positions) suggests that steric hindrance introduced by the affibody moiety disrupts proper nanocompartment assembly.

**A**

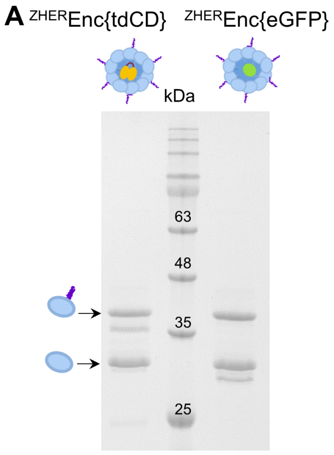

**B**

| | Shell proteins | |
|---|---|---|
| | $^{ZHER}$Enc | Enc |
| $^{ZHER}$Enc{tdCD} | 21.0 ± 0.8 | 39.0 ± 0.8 |
| $^{ZHER}$Enc{eGFP} | 22.2 ± 0.4 | 37.8 ± 0.4 |

**Figure S4**: Densitometric analysis of shell-modified encapsulin constructs. A) Representative SDS-PAGE gel of $^{ZHER}$Enc{tdCD} and $^{ZHER}$Enc{eGFP}. B) Densitometry analysis of the encapsulin shell composition, which consists of 60 identical subunits, a mixture of affibody-modified encapsulin ($^{ZHER}$Enc) and wild-type encapsulin (Enc). The table presents the calculated number of each subunit type per shell for $^{ZHER}$Enc{tdCD} and $^{ZHER}$Enc{eGFP} constructs.

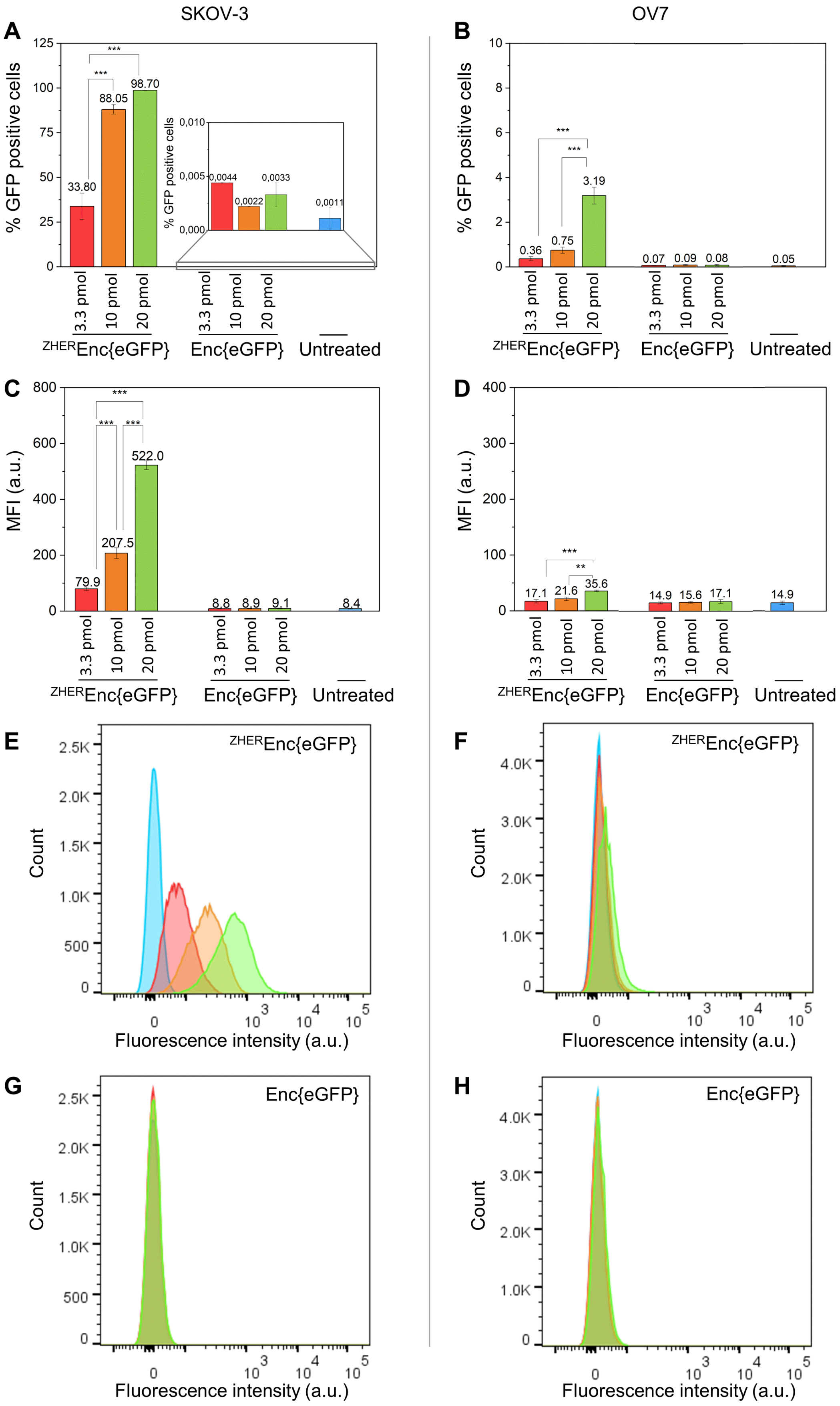

SKOV-3
OV7
A
B
C
D
E
F
G
H
% GFP positive cells
MFI (a.u.)
3.3 pmol
10 pmol
20 pmol
ZHEREnc{eGFP}
Enc{eGFP}
Untreated
33.80
88.05
98.70
0,0044
0,0022
0,0033
0,0011
0.36
0.75
3.19
0.07
0.09
0.08
0.05
79.9
207.5
522.0
8.8
8.9
9.1
8.4
17.1
21.6
35.6
14.9
15.6
17.1
14.9
Count
Fluorescence intensity (a.u.)

**Figure S5**: Flow cytometry analysis of $^{ZHER}$Enc{eGFP} and Enc{eGFP} binding to HER2 on the surface of SKOV-3 and OV7 cells. Cells were treated with $^{ZHER}$Enc{eGFP} or Enc{eGFP} at 3.3 pmol (red), 10 pmol (orange), or 20 pmol (green), and compared to untreated controls (blue). (A, B) Percentage of GFP-positive SKOV-3 (A) and OV7 (B) cells. (C, D) Mean fluorescence intensity (MFI) of SKOV-3 (C) and OV7 (D) cells. (E–H) Representative flow cytometry histograms showing fluorescence intensity distribution in SKOV-3 (E, G) and OV7 (F, H) cells treated with $^{ZHER}$Enc{eGFP} (E, F) or Enc{eGFP} (G, H). Statistical significance: * $p \leq 0.05$, ** $p \leq 0.01$, *** $p \leq 0.001$.

**References**

**1** Eigenbrot C, Ultsch M, Dubnovitsky A, Abrahmsén L, Härd T. Structural basis for high-affinity HER2 receptor binding by an engineered protein. *Proc Natl Acad Sci U S A*. 2010;107(34):15039-15044. doi:10.1073/pnas.1005025107.
